# Dissection of a sensorimotor circuit that regulates aversion to odors and pathogenic bacteria in *C. elegans* by whole-brain simulation

**DOI:** 10.1101/2022.04.21.489073

**Authors:** Adam Filipowicz, Jonathan Lalsiamthara, Alejandro Aballay

## Abstract

Altering behavior to reduce pathogen exposure is a key line of defense against pathogen attack for nearly all animals. The use of *Caenorhabditis elegans* bacterial infection models have allowed for many insights into the molecular mechanisms of behavioral immunity. However, the neural circuitry between chemosensory neurons that sense pathogenic bacterial cues and the motor neurons responsible for avoidance-associated locomotion remains unknown. We found that backward locomotion was a component of learned pathogen avoidance, as animals pre-exposed to *Pseudomonas aeruginosa* or *Enterococcus faecalis* showed reflexive aversion to drops of the bacteria, requiring ASI, AWB, and AWC neurons and ASE, AWB, and AWC neurons, respectively. This response also involved intestinal distention and, for *E. faecalis*, required expression of TRPM channels in the intestine and excretory system. Using whole-brain simulation and functional assays, we uncovered a sensorimotor circuit governing learned reflexive aversion. This behavior is controlled by a four-layer neural circuit composed of olfactory neurons, interneurons, and motor neurons that control backward locomotion crucial for learned reflexive aversion to pathogenic bacteria, learned avoidance, and a repulsive odor. The discovery of a complete sensorimotor circuit for reflexive aversion demonstrates the utility of using the *C. elegans* connectome and computational modeling in uncovering new neuronal regulators of behavior.

## Introduction

To reduce exposure to pathogens, organisms throughout the animal kingdom, including humans, engage in behavioral immune activities (1–4). Mechanistic insights into behavioral immunity are therefore crucial in furthering our understanding of human health and disease. Numerous studies have shown that *Caenorhabditis elegans* exhibits behavioral immunity in the form of pathogen avoidance strategies that improve its survival (4–9). These behaviors range from aversive reflexes to learned avoidance. Studies using pathogenic bacteria such as *Pseudomonas aeruginosa* and *Enterococcus faecalis*, for example, have shown that these bacteria are initially attractive to *C. elegans*, and it is only after hours-long exposure to the pathogens that the valence of the bacteria switches from attractive to aversive (9, 10). This is in contrast to the *C. elegans* avoidance of the toxins produced by *Streptomyces*, which takes place in a matter of seconds (7).

Taking advantage of the genetic tractability of *C. elegans*, the neuronal and molecular mechanisms of these behaviors have begun to be made clear. For example, the response to *Streptomyces* requires the G-protein-coupled receptor (GPCR) SRB-6, which is expressed in five different chemosensory neurons: ASH, ADL, ADF, PHA, and PHB (7). ASH, in particular, was shown to respond to both *Streptomyces* in an *srb-6-*dependent manner, though whether loss of ASH function leads to an abrogated behavioral response remains unknown. The avoidance of *P. aeruginosa* and *E. faecalis*, on the other hand, seems to involve a variety of bacterial cues sensed by various chemosensory neurons. These include bacterial metabolite sensation by DAF-7-expressing ASI and ASJ chemosensory neurons (11, 12), olfactory preference establishment by AWB and AWC olfactory neurons and modulation by serotonergic ADF neurons and RIA interneurons (9, 13), oxygen sensation by NPR-1 expressing AQR, PQR, and URX neurons (8, 14, 15), nitric oxide sensation by ASJ neurons (16), and BAG neuron-mediated sensation of carbon dioxide (17).

While the chemosensory neurons responsible for some forms of pathogen avoidance have been at least partially worked out, the complete neural circuitry required to coordinate these behaviors remains unknown. Signals from the sensory neurons described above likely converge on downstream interneurons. Activation of ASI neurons, for example, results in the release of the insulin-like-peptide INS-6, inhibiting *ins-7* expression in URX and subsequent upregulation of DAF-2 activity in RIA interneurons (18). These same interneurons are involved in the modulation of olfactory preference (13), and thus may be important integrators of distinct bacterial cues triggering avoidance behaviors.

Any chemosensory signal must eventually be propagated down not only to interneurons but also to the motor neurons required for avoidance-associated locomotion (19–21). The circuitry linking these neurons together is unknown, but the complete connectome of *C. elegans* (22) can be used to dissect the mechanisms involved in translating the detection of pathogenic cues into physical avoidance. We discovered that backward locomotion is a crucial component of pathogen avoidance, as animals trained on either *P. aeruginosa* or *E. faecalis* display reflexive aversion to these pathogens. To elucidate the reflexive aversion circuitry, we used simulations of the *C. elegans* nervous system, which have been shown to be useful in studying behaviorally relevant neural activity (23, 24). Using one such simulation platform, the *C. elegans* Neural Interactome (24), we investigated the neural patterns resulting from stimulation of the chemosensory neurons known to be involved in different pathogen avoidance behaviors. We found that oscillations in motor neurons critical for backward locomotion could be induced by AWB stimulation. AUA and RMG interneurons electrically coupled to AWB neurons also showed high activity upon AWB stimulation, and *in silico* ablation of these neurons resulted in the loss of motor neuron oscillations. Genetic ablation of these neurons demonstrated their involvement not only in pathogen avoidance, but 2-nonanone aversion as well. The olfactory neuron AWB, electrically synapses onto AUA and RMG interneurons, which themselves synapse onto motor command interneurons to control backward locomotion motor neurons, thus representing a novel sensorimotor circuit for pathogen and repulsive odor aversion.

## Results

### Intestinal infection by *P. aeruginosa* or *E. faecalis* induces a learned reflexive aversion requiring multiple chemosensory neurons

To investigate the neural circuitry governing the translation of pathogen chemosensory cues into the motor neuron activity necessary for avoiding said cues, we directly tested avoidance locomotion by making use of an assay that would allow us to quickly assess individual neuron requirements for reflexive aversion both before and after exposure to a pathogen (Figure 1A). The pathogenic bacteria *P. aeruginosa* and *E. faecalis* are initially attractive to *C. elegans* and only induce an avoidance response after many hours of exposure (4, 9, 10, 13). This learning process involves the association of infection and subsequent physiological responses, including intestinal distention, engagement of RNAi pathways, and immune activation, with bacterial cues, resulting in avoidance of the bacteria (10, 12, 25). Using the reflexive aversion assay, we found that naïve animals do not respond to drops of *P. aeruginosa* (Figure 1B) or *E. faecalis* (Figure 1C); however, animals exposed to bacteria prior to testing showed reflexive aversion to the same bacteria. The response to *P. aeruginosa* required ASI, AWB, and AWC neurons, while the response to *E. faecalis* required ASE, AWB, and AWC neurons (Figure 1B and 1C). These results indicate that intersecting neural circuits are required for learned reflexive aversion against different pathogens.

**Fig. 1.**
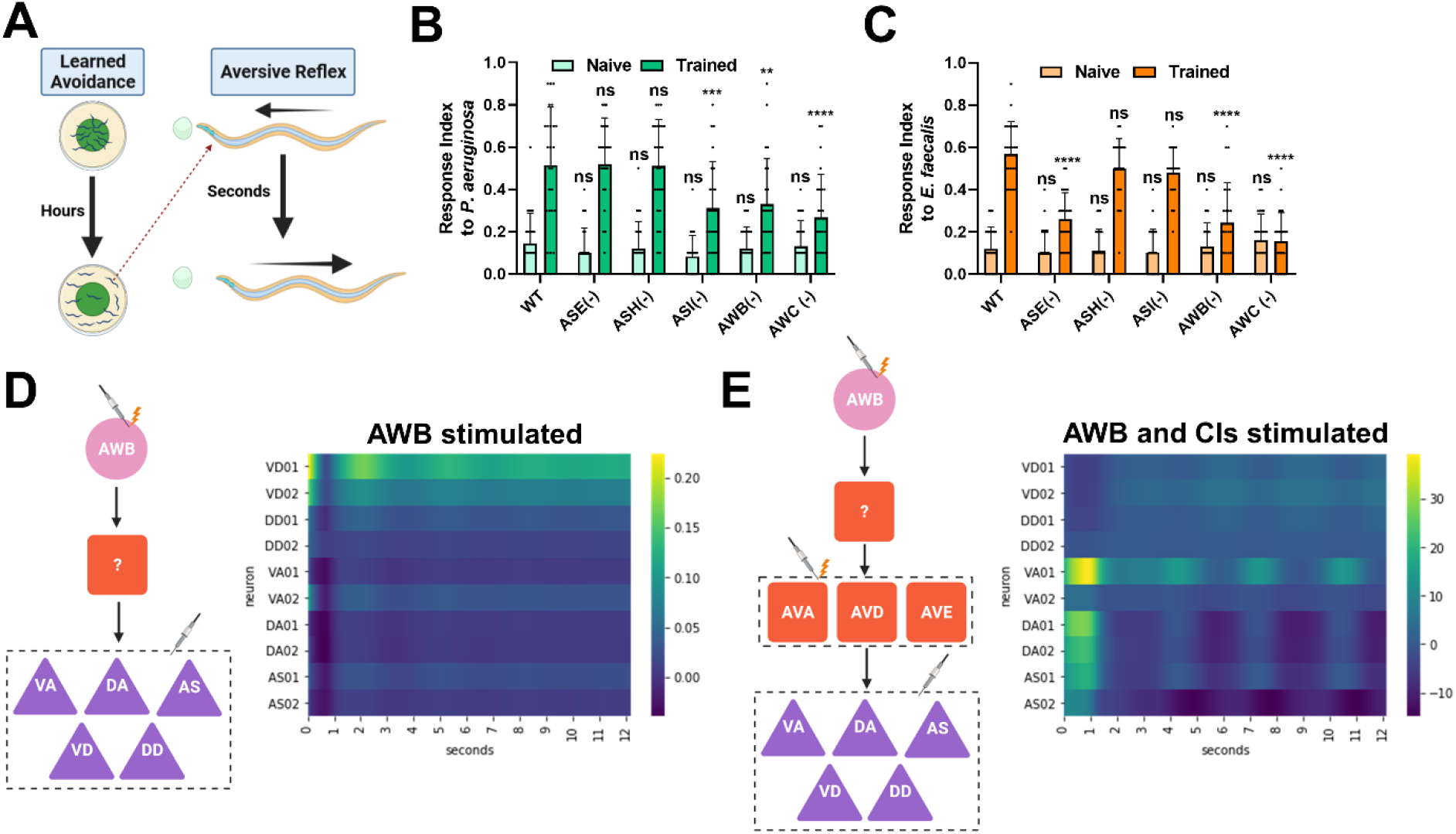
Intestinal infection induces a learned reflexive aversion requiring multiple chemosensory neurons. (A) Diagram of the assays to determine learned avoidance (left) and naïve reflexive aversion (right). Details of assays can be found in the Materials and Methods. Briefly, for learned avoidance, animals are placed on a lawn of bacteria and allowed to roam freely for several hours (4 hr for *E. faecalis*, 24 hr for *P. aeruginosa*) before the number of animals inside and outside the bacterial lawn is counted. For reflexive aversion, a drop of bacterial culture is placed in front of a forward-moving animal, and a response is recorded if the animal initiates backward locomotion upon encountering the dried drop. Trained reflexive aversion involves taking from the avoidance plates and placing them onto a plate with no bacteria. Then, the reflexive aversion assay is carried out. (B) Response index to *P. aeruginosa* for both naïve (light green) and trained (dark green) animals with either no neurons ablated (N2, WT) or ASE (PR680), ASH (JN1713), ASI (PY7505), AWB (JN1715), or AWC (PY7502) neurons ablated. (C) Response index to *E. faecalis* for the same groups as in **B**. For both **B** and **C**, two-way ANOVA with subsequent comparison to naïve or trained WT groups was performed. Error bars depict standard deviation. N = 25 (individual dots) for all groups. (D) Schematic of the sensorimotor circuit and protocol used in the Neural Interactome (left). AWB neurons were stimulated at 5.0 nA and the activity of VA, DA, AS, VD, and DD motor neurons (dashed-outline) was recorded. There is no direct connection between AWB neurons and the motor neurons, so an unknown interneuron must complete the circuit (question mark). Pink circle = sensory neuron; red square = interneuron; purple triangle = motor neuron. The recorded activity of the motor neurons is presented as a heatmap (right), with rows representing individual neuronal activity over time. The first two neurons of each motor neuron class were chosen for ease of visualization. (E) An updated schematic of the stimulation protocol (left). AVA, AVD, and AVE command interneurons (CIs) were stimulated at 0.9 nA along with the 5.0 nA stimulation of AWB neurons. This resulted in oscillations in the motor neurons (right). There is no direct connection between AWB neurons and the CIs, so another interneuron must complete the circuit (question mark).

### Nervous system simulation predicts that AUA and RMG neurons are in the AWB neuron-mediated learned reflexive aversion circuit

To uncover the overall neural circuitry that integrates bacterial-related cues that result in learned reflexive aversion, we used the *C. elegans* Neural Interactome, a simple, user-friendly simulation of the *C. elegans* nervous system that allows for stimulation and ablation of individual neurons and outputs total network activity (24). The simulation takes advantage of the complete connectome of *C. elegans* and its neural dynamics, and was previously shown to assist in the study of neural response patterns associated with locomotion and external stimuli such as nose touch. This makes the Neural Interactome a potentially useful tool in studying other aversive behaviors, including pathogen avoidance. To assess the neural activity of reflexive aversion, we measured oscillations in VA, DA, VD, DD, and AS motor neurons as a readout for backward locomotion, as it has previously been shown that these neurons are active during backward locomotion and function as oscillators (19, 20, 24, 26). Because AWB neurons are at the intersection of the circuits required for learned reflexive aversion against *P. aeruginosa* and *E. faecalis*, we first stimulated AWB neurons. However, we found no oscillation in the aforementioned motor neurons (Figure 1D), indicating that additional neurons must participate in the circuit. Previous studies indicated that the command interneurons (CIs) AVA, AVD, and AVE necessary reflexive aversion to nose touch (21, 24). Thus, we used the Neural Interactome to activate AWB together with CIs and found oscillatory activity in the VA, DA, VD, DD, and AS motor neurons (Figure 1E). Stimulation of AWC and ASI neurons led to no oscillations in the motor neurons, while stimulation of ASE neurons led to very weak oscillations (Figure S1). This indicates that AWB neurons are the primary mediator of reflexive aversion, while AWC, ASI, and ASE play some other roles in the learning process.

To confirm that motor neuron oscillations are a good readout for aversion behaviors, we examined a behavior that has a known circuit correlate: reflexive aversion to the common laboratory detergent sodium dodecyl sulfate (SDS; (27)). We confirmed that this behavior requires the chemosensory neuron ASH, as animals lacking ASH neurons responded less strongly to SDS compared to wild-type animals (Figure S2A). Aversion to SDS requires the command interneurons AVA, AVD, and AVE, and the motor neurons VA, DA, VD, DD, and AS (19, 20, 27). The connections between these neurons resolve as a three-layer circuit (Figure S2B). Stimulation of ASH, AVA, AVD, and AVE neurons within the Neural Interactome resulted in oscillatory activity in VA, DA, DD, and AS motor neurons, indicating that oscillations in these neurons were a suitable readout for aversion behaviors (Figure S2C). Furthermore, this same circuitry is required for the response to dodecanoic acid (Figure S2A). Dodecanoic acid is a toxin produced by the pathogenic bacteria *Streptomyces* (7), thus showing the simulation’s ability to uncover circuits of pathogen avoidance.

AWB neurons have no direct connection to the motor command AVA, AVD, and AVE interneurons (Figure 1E). Thus, another layer of interneurons is likely involved in the circuit. Looking at the activity of all the neurons in the interactome model after AWB stimulation, the AUA and RMG neurons stood out as having high oscillatory activity (Figures 2A and B). According to the connectome of *C. elegans*, AUA and RMG form electrical synapses with AWB, and chemically synapse onto the AVA, AVD, and AVE neurons (Figure 2C; (22)). This places them in a prime position to be regulators of the circuit for learned reflexive aversion. Indeed, *in silico* ablation of either AUA or RMG neurons resulted in the loss of oscillatory activity in VA, DA, VD, DD, and AS motor neurons upon AWB, AVA, AVD, and AVE stimulation (Figure 2D and E). Other neurons, such as AIB, AVB, and SMB, also bridge the AWB and motor command interneurons, but *in silico* ablation of these neurons left the motor neuron oscillations intact (Figure S3).

**Fig. 2.**
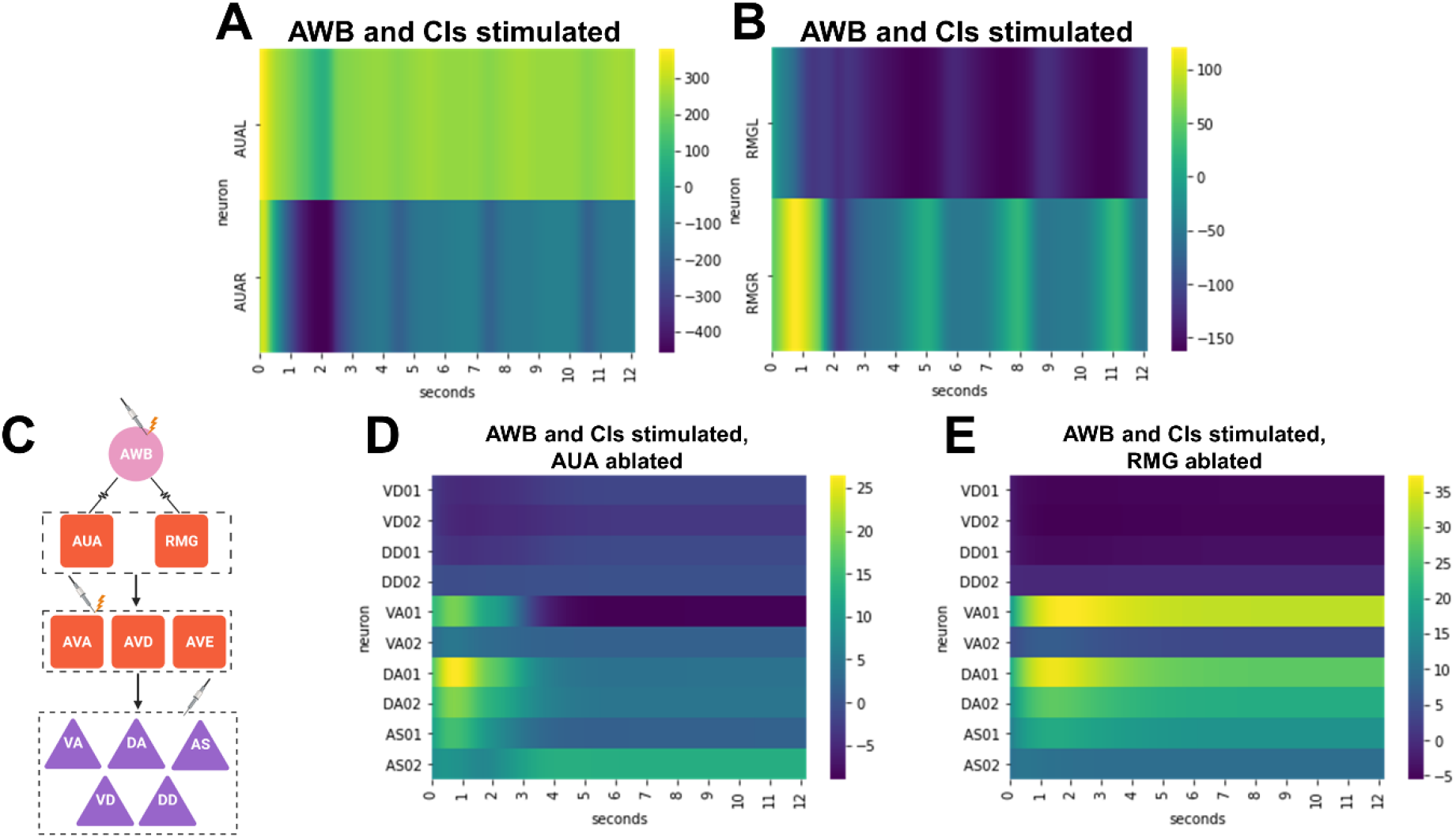
Nervous system simulation predicts that AUA and RMG neurons are in the AWB neuron-mediated learned reflexive aversion circuit. (A) Activity of AUA neurons (rows) upon 5.0 nA stimulation of AWB neurons and 0.9 nA stimulation of the CIs in the Neural Interactome. (B) Activity of RMG neurons (rows) upon 5.0 nA stimulation of AWB neurons and 0.9 nA stimulation of the CIs in the Neural Interactome. (C) Updated circuit diagram and stimulation protocol schematic showing the connections between AWB, AUA, RMG, AVA, AVD, AVE, VA, DA, AS, VD, and DD neurons. The first neuron of each motor neuron class was chosen for ease of visualization. Arrows represent chemical synapses, while jagged lines represent electrical synapses. (D) Activity of motor neurons (rows) upon 5.0 nA stimulation of AWB neurons and 0.9 nA stimulation of the CIs with AUA neurons ablated in the Neural Interactome. (E) Activity of motor neurons (rows) upon 5.0 nA stimulation of AWB neurons and 0.9 nA stimulation of the CIs with RMG neurons ablated in the Neural Interactome.

### A four-layer circuit underlies pathogen avoidance behaviors and aversion to 2-nonanone

To test whether AUA and RMG neurons affected learned reflexive aversion, we genetically ablated AUA and RMG neurons (Figure S4) and tested their response to drops of *P. aeruginosa* and *E. faecalis*. Ablation of either AUA or RMG left naïve responses to both bacteria intact, while trained responses showed a significant decrease (Figures 3A and B). We also measured the occupancy index of animals lacking either AUA or RMG for both *P. aeruginosa* and *E. faecalis* and found that loss of either neuron resulted in increased lawn occupancy (Figure 3C and D). This indicates that AUA and RMG neurons contribute to both learned reflexive aversion and learned pathogen avoidance of *P. aeruginosa* and *E. faecalis*.

**Fig. 3.**
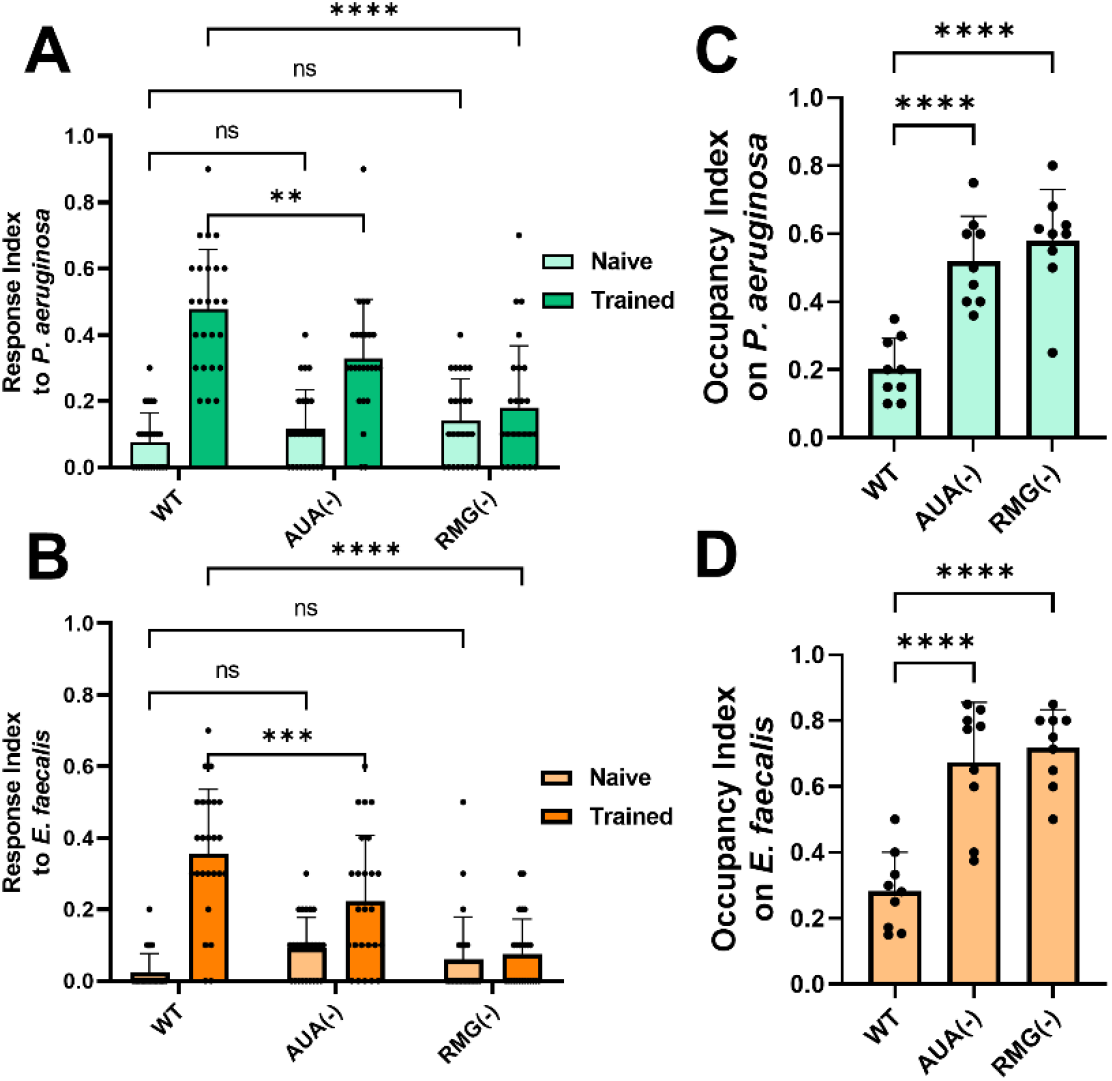
A four-layer circuit underlies pathogen avoidance behaviors. (A) Response index to *P. aeruginosa* for both naïve (light green) and trained (dark green) animals with either no neurons ablated (WT) or AUA (AY178) or RMG (AY179) neurons ablated. (B) Response index to *E. faecalis* for the same groups as in **A**. For both **A** and **B**, two-way ANOVA with subsequent comparison to naïve or trained WT groups was performed. Error bars depict standard deviation. N = 25 (individual dots) for all groups. (C) Occupancy index for *P. aeruginosa* after 24 hours for animals with no neurons ablated (WT) or AUA or RMG neurons ablated. (D) Occupancy index for *E. faecalis* after 4 hours for the same groups as in **C**. For both **C** and **D**, one-way ANOVA with subsequent comparison to the WT group was performed. Error bars depict standard deviation. N = 9 (individual dots) for all groups.

We wondered whether the learned reflexive aversion response was specific to the bacteria used to train the animals or if it was a general result of bacterial infection. We cross-tested animals, training them on *E. faecalis* and then testing them with drops of *P. aeruginosa* and vice versa, and found that animals did not show any reflexive aversion (Figure 4A and 4B). However, if *E. faecalis* lawns were paired with the odor of *P. aeruginosa* during training, achieved by placing an agar plug of *P. aeruginosa* on the lid of the training plates, animals showed reflexive aversion (Figure 4A). The same was true for the opposite pairing (Figure 4B), suggesting that underlying learned reflexive aversion is a process whereby animals associate the odors of the bacteria with infection.

**Fig. 4.**
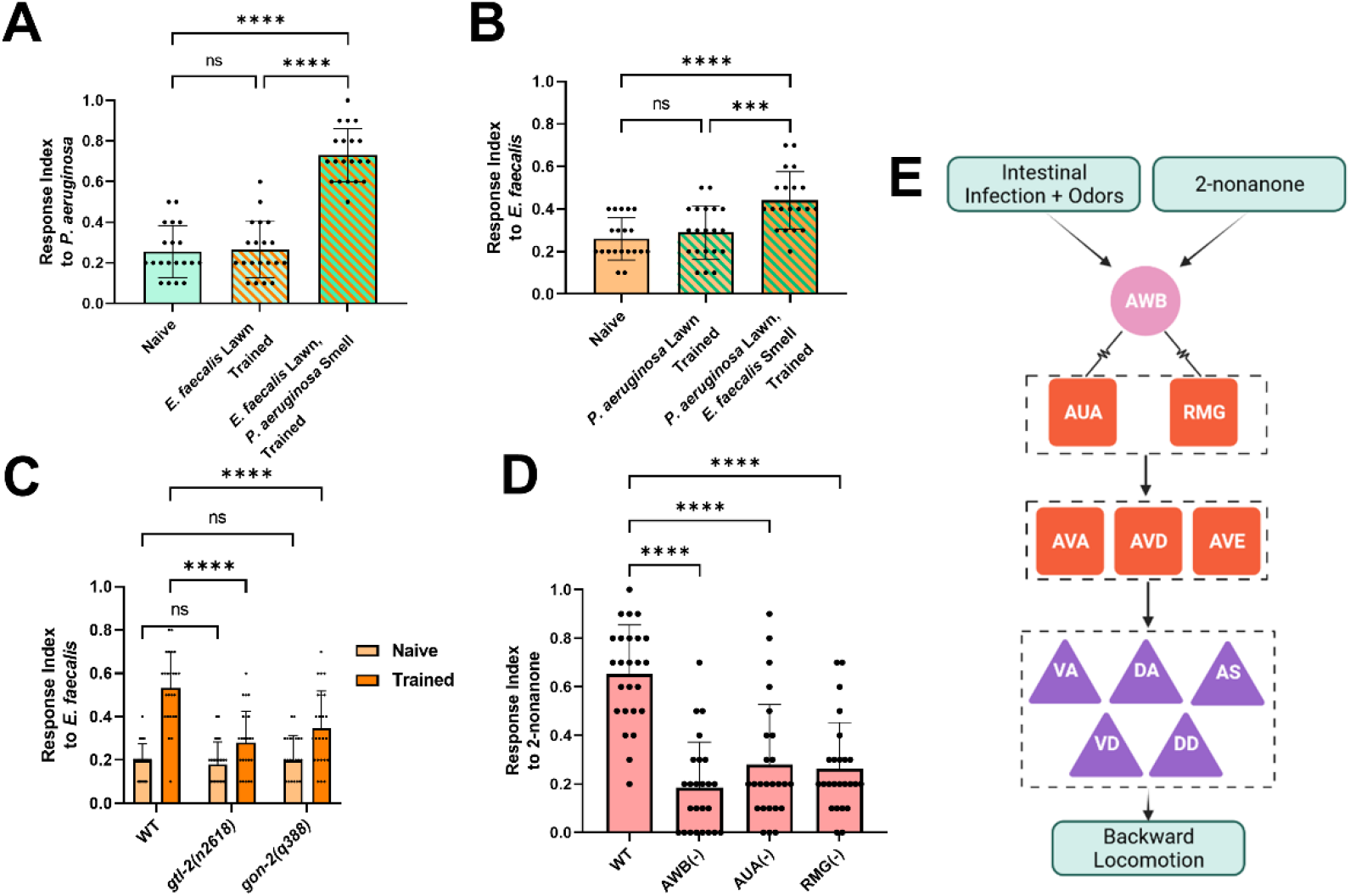
The sensorimotor circuit involves olfaction and regulates aversion to 2-nonanone. (A) Response index to *P. aeruginosa* for wild-type animals either naïve to *P. aeruginosa*, trained on *E. faecalis* lawns, or trained on *E. faecalis* lawns with a lawn of *P. aeruginosa* on the lid, inaccessible to the animals. (B) Response index to *E. faecalis* for wild-type animals either naïve to *E. faecalis*, trained on *P. aeruginosa*, or trained on *P. aeruginosa* lawns with a lawn of *E. faecalis* on the lid. For both **A** and **B**, one-way ANOVA with subsequent comparisons between all groups was performed. Error bars depict standard deviation. N = 25 (individual dots) for all groups. (C) Naïve and trained response index to *E. faecalis* for wild-type (WT) and either *gtl-2* or *gon-2* loss-of-function mutants. Two-way ANOVA with subsequent comparison to the WT groups was performed. Error bars depict standard deviation. N = 25 (individual dots) for all groups. (D) Response index to 2-nonanone (1:10) for animals with no neuronal ablation (WT) or AWB, AUA, or RMG neurons ablated. One-way ANOVA with subsequent comparison to the WT group was performed. Error bars depict standard deviation. N = 25 (individual dots) for all groups. (E) Diagram of the odor-aversion sensorimotor circuit. AWB neurons sense an olfactory cue either from pathogenic bacteria or other repulsive odorants such as 2-nonanone. They pass this signal to AUA and RMG neurons via electrical synapses. AUA and RMG neurons form chemical synapses with AVA, AVD, and AVE command interneurons, which synapse with the motor neurons important for backward locomotion, VA, DA, VD, DD, and AS neurons. These neurons execute the backward locomotion necessary for odor avoidance.

Because we noticed a significant variation in the reflexive aversion of trained animals, we wondered whether varying levels of intestinal distention, which can trigger learned avoidance (10, 25), accounted for the variation in the responses. Intestinal distention on *P. aeruginosa*, measured by either PA14-GFP signal in the intestinal lumen or intestine diameter, correlated with the trained response index to a weak, but significant, amount (Figure S5A and B). However, intestinal distention on *E. faecalis* showed no such correlation, perhaps due to the fact that distention was much more prevalent and severe in animals exposed to *E. faecalis* (Figure S5C and D). The association of intestinal distention and *E. faecalis* cues requires the TRPM channels GON-2 and GTL-2 (10). We found that these channels were also required for the learned reflexive aversion to *E. faecalis*, as animals with loss-of-function mutations in the genes encoding for GON-2 and GTL-2 displayed mitigated responses to *E. faecalis* after training (Figure 4C). Although intestinal distention may not explain the variation in aversion observed, this data suggests that the association of intestinal distention and bacterial cues is required for the trained reflexive aversion to at least *E. faecalis*, indicating that the mechanisms of learned reflexive aversion and learned avoidance share similar connections.

Finally, we wondered whether this circuit was specific to the pathogen response or if it governed a general reflexive aversion to repulsive odors. We found that animals lacking AWB neurons showed reduced reflexive aversion to 2-nonanone, a volatile organic compound that is known to be repulsive to *C. elegans* (Figure 4D). This confirms previous reports that AWB is the primary senor of 2-nonanone (28, 29). Importantly, animals lacking either AUA or RMG also showed reduced reflexive aversion to 2-nonanone (Figure 4D). Altogether, as illustrated in Figure 4E, these results indicate that learned reflexive aversion involves the formation of an association between intestinal infection and bacterial odors, with the behavior executed by a four-layer neural circuit composed of AWB olfactory neurons electrically synapsed to AUA and RMG interneurons, which themselves are connected chemically to the motor-command interneurons AVA, AVD, and AVE. These interneurons control the motor neurons, VA, DA, VD, DD, and AS, which execute backward locomotion. Backward locomotion and the underlying circuit seem to be crucial for learned reflexive aversion to *P. aeruginosa* and *E. faecalis*, learned avoidance of lawns of these pathogens, and aversion to a repulsive volatile organic compound, 2-nonanone, in *C. elegans*. Taken together, the results implicate the sensorimotor circuit in the control of an innate reflexive aversion to a repulsive odor and a learned aversion to pathogenic bacteria.

## Discussion

The discovery of a sensorimotor circuit involved in learned reflexive aversion to the pathogenic bacteria *P. aeruginosa* and *E. faecalis* and naïve aversion to 2-nonanone demonstrates the utility of the *C. elegans* Neural Interactome as a tool for hypothesis generation. AUA and RMG neurons had not previously been implicated in either pathogen avoidance or aversion to 2-nonanone. Interestingly, both had been implicated in the regulation of social feeding behavior (30, 31). AUA is a synaptic target of URX, and expression of *npr-1* in *npr-1(ad609)* mutants in AQR, PQR, URX, and AUA neurons results in suppression of aggregation and bordering behaviors (30). RMG is at the center of a gap junction hub-and-spoke circuit, connected to many sensory neurons, including ASK, URX, ASH, ADL, and AWB (31). High activity in RMG is essential for all aspects of social behavior, including aggregation, bordering, and, with input from ASK neurons, attraction to hermaphrodite pheromones. The innexin gene *unc-9* was shown to be required in RMG neurons to drive social behavior (32). As AWB and AUA neurons also express *unc-9* (33), it is possible that *unc-9*-based gap junctions are also required for AWB-mediated pathogen and odor avoidance behaviors.

A crucial step to study sensorimotor circuits necessary for learned pathogen avoidance was to find a behavior that matched the seconds-long timescale necessary to perform simulations. Learned pathogen avoidance takes hours and was therefore not a suitable behavior. The discovery that *C. elegans* avoids drops of *P. aeruginosa* and *E. faecalis* within seconds, following hours-long pre-exposure on bacterial lawns, allowed us to match the behavior and the simulated data from the Interactome on similar timescales. Ultimately the neurons involved in learned reflexive aversion and learned pathogen avoidance were the same for both *P. aeruginosa* and *E. faecalis*, with AUA and RMG neurons required for both (Figure 3). Thus, we speculate that learned reflexive aversion to pathogenic bacteria is a crucial component of the general learned pathogen avoidance behavior. In general avoidance, the reflex to avoid pathogenic bacteria is most likely countered by attraction to the bacteria as a food source. This attraction is likely driven by AWC neurons, which are known to shape the olfactory response to pathogenic bacteria and food odors (9, 34). We found that AWC neurons were required for the learned reflexive aversion response, further implicating them in the associative learning process required for avoidance of *P. aeruginosa* and *E. faecalis* (Figures 1B and C). The cross-training experiments performed with the two pathogens make clear that this process is driven by olfaction (Figures 4A and B). Exactly how this learning takes place, and what the changes in the neural dynamics are that allow for a shift from attraction to aversion over time remains unclear, though modulation via serotonin likely plays a role (9, 13). ASI and ASE neurons also seem to play a role in the learned aversion to *P. aeruginosa* and *E. faecalis*, respectively (Figure 2B and C). The simulated data suggest that such a role is not through the engagement of the backward-movement motor neuron circuitry, as stimulation of AWC, ASE, or ASI neurons does not lead to strong oscillations in the motor neurons (Figure S1). This is in contrast to the stimulation of AWB neurons, which induces motor neuron oscillations (Figure 1E). What bacterial cue actually stimulates the AWB neurons remains unknown, though it is likely a bacterial odor (or blend of odors). This could be an initially attractive odor that becomes aversive with bacterial infection, or an innately aversive odor such as 1-undecene (6).

While the Interactome allowed us to accurately predict which neurons would be necessary for pathogen avoidance and 2-nonanone aversion, the actual neural dynamics at play in these behaviors may be quite different. An important next step would be to experimentally uncover the dynamics at play and compare them to the Interactome model. This would allow for further refinement of the model, and would be possible using tools such as NeuroPAL, which allows for neuronal identification of all *C. elegans* neurons via a multicolor fluorescence map, and genetically encoded calcium indicators (35). Further combinations of experimental manipulation and computational modeling will be crucial in deepening our understanding of behaviorally relevant neural circuits.

## Materials and Methods

### Bacterial strains

The following bacterial strains were used: *Enterococcus faecalis* OG1RF, *E. faecalis* OG1RF-GFP, *Escherichia coli* OP50, *Pseudomonas aeruginosa* PA14, and *P. aeruginosa* PA14-GFP. *E. coli* and *P. aeruginosa* bacterial strains were grown in Luria-Bertani (LB) broth at 37°C, while the *E. faecalis* was grown in brain-heart infusion (BHI) broth at 37°C.

#### *C. elegans* strains and maintenance

The following *C. elegans* strains used in this study were obtained from the CGC: N2 (WT strain), CZ9957 *gtl-2(n2618)*, EJ1158 *gon-2(q388)*, JN1713 peIs1713 [*sra-6p::mCasp-1 + unc-122p::mCherry*], JN1715 peIs1715 [*str-1p::mCasp-1 + unc-122p::GFP*], NY2078 ynIs78 [*flp-8p::GFP*], NY2087 ynIs87 [*flp-21p::GFP*], PR680 *che-1(p680)*, PY7502 oyIs85 [*ceh-36p::TU#813 + ceh-36p::TU#814 + srtx-1p::GFP + unc-122p:DsRed*], and PY7505 oyIs84 [*gpa-4p::TU#813 + gcy-27p::TU#814 + gcy-27p::GFP + unc-122p::DsRed*]. The following transgenic strains were generated for this study using standard microinjection protocols (10): AY178 ynIs78 [*flp-8p::GFP*]; *flp-8p::ced-3 (p15)::nz + flp-32::cz::ced-3 (p17) + unc-122p::rfp* and AY179 ynIs87 [*flp-21p::GFP*]; *flp-21p::ced-3 (p15)::nz + ncs-1p::cz::ced-3 (p17) + unc-122p::rfp. C. elegans* hermaphrodites were maintained on *E. coli* OP50 at 20°C.

### Construction of neuronal ablation strains

The transgenic strains AY178 (AUA ablation) and AY179 (RMG ablation) were generated by first constructing pJL12 (pPD95.75 *flp-8p::ced-3 (p15)::nz*), pJL13 (pPD95.75 *flp-32p::cz::ced-3 (p17)*), pJL14 (pPD95.75 *flp-21p::ced-3 (p15)::nz*), and pJL15 (pPD95.75 *ncs-1p:: cz::ced-3 (p17)*) plasmids. The pJL12 plasmid was constructed by cloning a 2019 bp upstream-promoter region of the *flp-8* gene into a *ced-3 (p15)::nz* backbone vector, via SphI-BamHI restriction sites. The pJL13 plasmid was constructed by cloning a 2085 bp upstream-promoter region of the *flp-32* gene into a cz::ced-3 (p17) backbone vector, via SphI-BamHI restriction sites. The pJL14 plasmid was constructed by cloning a 4109 bp upstream-promoter region of the *flp-21* gene into a *ced-3 (p15)::nz* backbone vector, via SphI-BamHI restriction sites. The pJL15 plasmid was constructed by cloning a 3116 bp upstream-promoter region of the *ncs-1* gene into a *cz::ced-3 (p17)* backbone vector, via SphI-BamHI restriction sites. recCaspase plasmids were a gift from Martin Chalfie, Addgene plasmids # 16080 and # 16081, Addgene, MA (36). A cocktail of pJL12 (10ng/μl), pJL13 (10ng/μl), co-injection marker *unc-122p::RFP* (50ng/μl) and empty vector PUC18 (50ng/μl) plasmids was co-injected into NY2078 animals to generate AY178 animals. A cocktail of pJL14 (10ng/μl), pJL15 (10ng/μl), co-injection marker *unc-122p::RFP* (50ng/μl) and empty vector PUC18 (50ng/μl) plasmids was co-injected into NY2087 animals to generate AY179 animals. Transgenic animals showing successful ablation of AUA or RMG neurons were selected and used for further assays. Plasmids were maintained as extrachromosomal arrays.

### Dry-drop assay

The dry-drop assay was carried out as previously described (7). Using a capillary, a drop of the appropriate stimulus was placed on a dry SK plate (for SDS, dodecanoic acid, 2-nonanone, and *P. aeruginosa*) or BHI plate (for *E. faecalis*) in front of a forward-moving animal. A response was counted if an animal initiated backward movement upon encountering the dried drop. Response Index = N_responses_/N_drops_. Ten drops were used per animal, and a total of 25 animals of each strain were used for each experiment. The stimuli were prepared as follows: 0.6mM SDS, 1mM dodecanoic acid, 1:10 2-nonanone, and a 5-6 hr liquid culture of either *P. aeruginosa* or *E. faecalis*.

### Lawn avoidance assays

Lawn avoidance assays were carried out as previously described (10). Briefly, bacterial cultures were grown by inoculating *P. aeruginosa* or *E. faecalis* colonies into 2 mL of either LB or BHI broth, respectively, and growing them for 5-6 hr on a shaker at 37°C. Then 20 μL of the culture was plated onto the center of 3.5-cm-diameter BHI or standard slow-killing (SK) plates (modified NGM agar plates [0.35% instead of 0.25% peptone]). The plates were then incubated overnight at 37°C. The plates were cooled to room temperature for at least 30 min before seeding with animals. Synchronized young gravid adult hermaphroditic animals grown on *E. coli* OP50 were washed with M9 buffer and transferred outside the bacterial lawns, and the number of animals on and off the lawns were counted at 24 hr for *P. aeruginosa* and 4 hr for *E. faecalis*. Three 3.5-cm-diameter plates were used per trial in every experiment. Occupancy index was calculated as N_on lawn_/N_total_.

### Aversive training

Training plates of 3.5-cm-diameter containing either *P. aeruginosa* on SK agar or *E. faecalis* on BHI agar were made as described above for avoidance assays. Young gravid adult hermaphroditic animals grown on *E. coli* OP50 were washed with M9 and transferred to the training plates and allowed to roam for 24 hr on *P. aeruginosa* and 4 hr on *E. faecalis*. They were then transferred to the appropriate assay plates. For cross-training with odors (Figure 2D and E), agar plugs with bacterial lawns were cut from growth plates and transferred to the lids of the training plates with the appropriate training bacteria, as previously described (10).

### Imaging and quantification

Fluorescence imaging was carried out as described previously (10). Briefly, the animals were anesthetized using an M9 salt solution containing 50 mM sodium azide and mounted onto 2% agar pads. The animals were then visualized using a Leica M165 FC fluorescence stereomicroscope. For quantification of intestinal lumen distention, brightfield images were acquired at 24 hr for *P. aeruginosa* and 4 hr for *E. faecalis* using the Leica LAS v4.6 software, and the diameter of the intestinal lumen was measured using ImageJ software. For quantification of fluorescent bacteria, fluorescent images were acquired using the Leica LAS v4.6 software, and ImageJ software was used to first draw a region of interest around the bacteria in the intestines of animals and then to measure fluorescence intensity in the region.

### Neural Interactome simulations and visualization

Simulations of the *C. elegans* nervous system were carried out using a local copy of the Neural Interactome, pulled from https://github.com/shlizee/C-elegans-Neural-Interactome. Simulations were run for at least 12 seconds and neuronal activity for each simulation was saved into a numpy file. Simulated data was loaded directly with the Python Numpy package into a Jupyter Notebook and converted to a Pandas DataFrame. Python scripts were written to visualize individual neuronal activity over time as rows on a heatmap and graphs were generated using the Matplotlib and Seaborn libraries. The first two neurons of the VD, DD, VA, DA, and AS motor neuron classes were chosen as representatives of each class to aid visualization.

### Statistical analysis

The statistical analysis was performed with Prism 9 (GraphPad). All bar graphs depict the mean of the population, with individual dots representing individual trials. Error bars represent the standard deviation of the population. For x-y correlation plots, individual dots represent measurements for individual animals, and the red line depicts a simple linear regression. All experiments were performed in triplicate on at least three separate days, resulting in the sample sizes listed in the figure legends. One- or two-way ANOVA with subsequent group comparisons were performed as indicated in the figure legends. In the figures, ns denotes not significant, and asterisks (*) denote statistical significance as follows: *≤0.05; **p≤0.01; ***p≤0.001; ****p≤0.0001, as compared with the appropriate controls.

## Acknowledgments

We thank the Garsin lab (McGovern Medical School) for providing the *E. faecalis* strains and the Ausubel lab (Harvard University) for providing the *P. aeruginosa* strains. We also thank members of the Vollum Institute, Neuroscience Graduate Program, and Graduate Researchers United at OHSU for their comments and support. Some strains used in this study were provided by the Caenorhabditis Genetics Center (CGC), which is funded by the NIH Office of Research Infrastructure Programs (P40 OD010440). Images were created using Microsoft PowerPoint, Biorender.com, and the Python programming language.

## Supplementary Figures

**Fig. S1.**
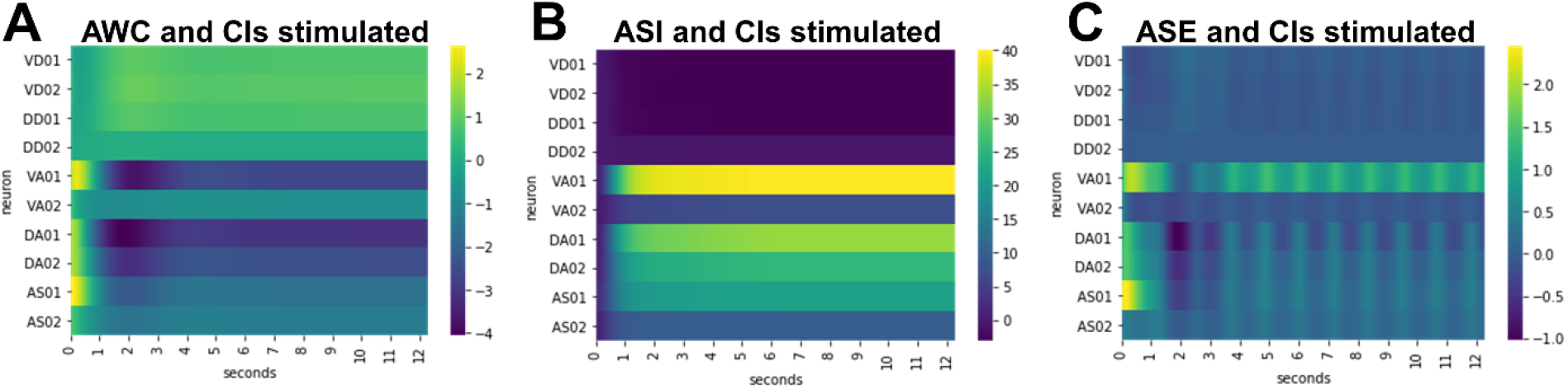
AWC, ASI, and ASE neurons do not lead to oscillations in motor neurons important for backward locomotion. Activity of VD, DD, VA, DA, and AS motor neurons in the Neural Interactome upon 0.9 nA stimulation of the CIs and 5.0 nA stimulation of AWC (A), ASI (B), and ASE (C).

**Fig. S2.**
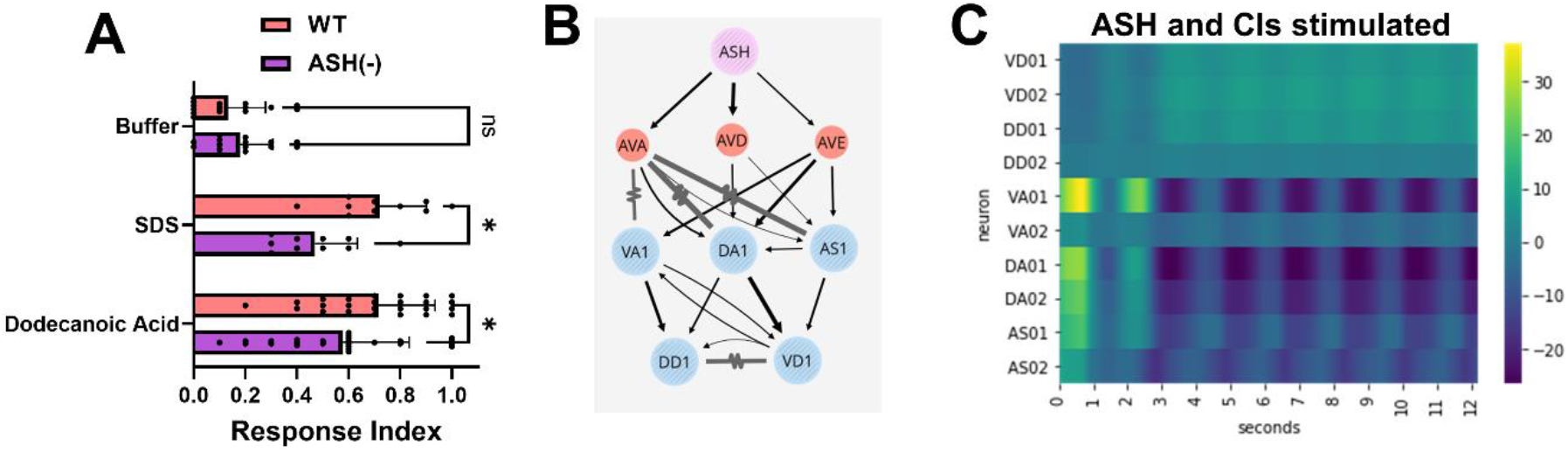
The response to SDS and dodecanoic acid are ASH mediated and can be modeled in the Neural Interactome. (A) Response index to buffer, 0.6 mM SDS, or 1mM dodecanoic acid for animals with no neurons ablated (WT, red) or ASH neurons ablated (ASH(-), purple). Two-way ANOVA with subsequent comparison to the WT groups was performed. Error bars depict standard deviation. N = 15 for buffer, 10 for SDS, and 25 for dodecanoic acid (individual dots). (B) Diagram of the circuit for reflexive aversion to SDS and dodecanoic acid. Arrows represent chemical synapses, while jagged lines represent electrical synapses. (C) Activity for the motor neurons VD, DD, VA, DA, and AS (rows) upon 5.0 nA stimulation of ASH neurons and 0.9 nA stimulation of the CIs in the Neural Interactome.

**Fig. S3.**
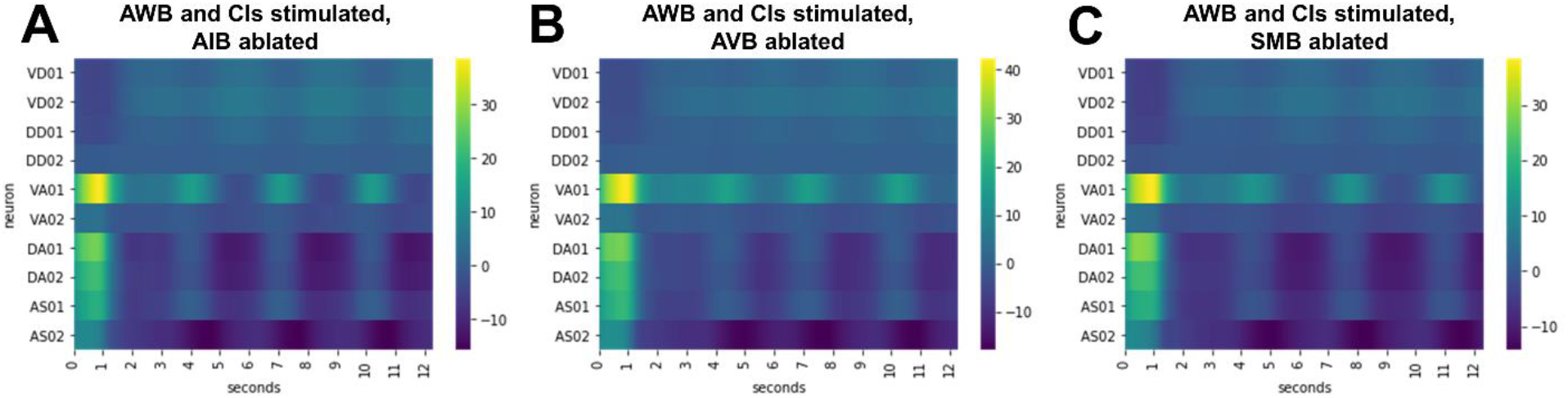
AIB, AVB, or SMB neuron ablation does not diminish oscillations in backward locomotion-associated motor neurons. Activity of motor neurons (rows) upon 5.0 nA stimulation of AWB neurons and 0.9 nA stimulation of the CIs with AIB (A), AVB (B), or SMB (C) neurons ablated in the Neural Interactome.

**Fig. S4.**
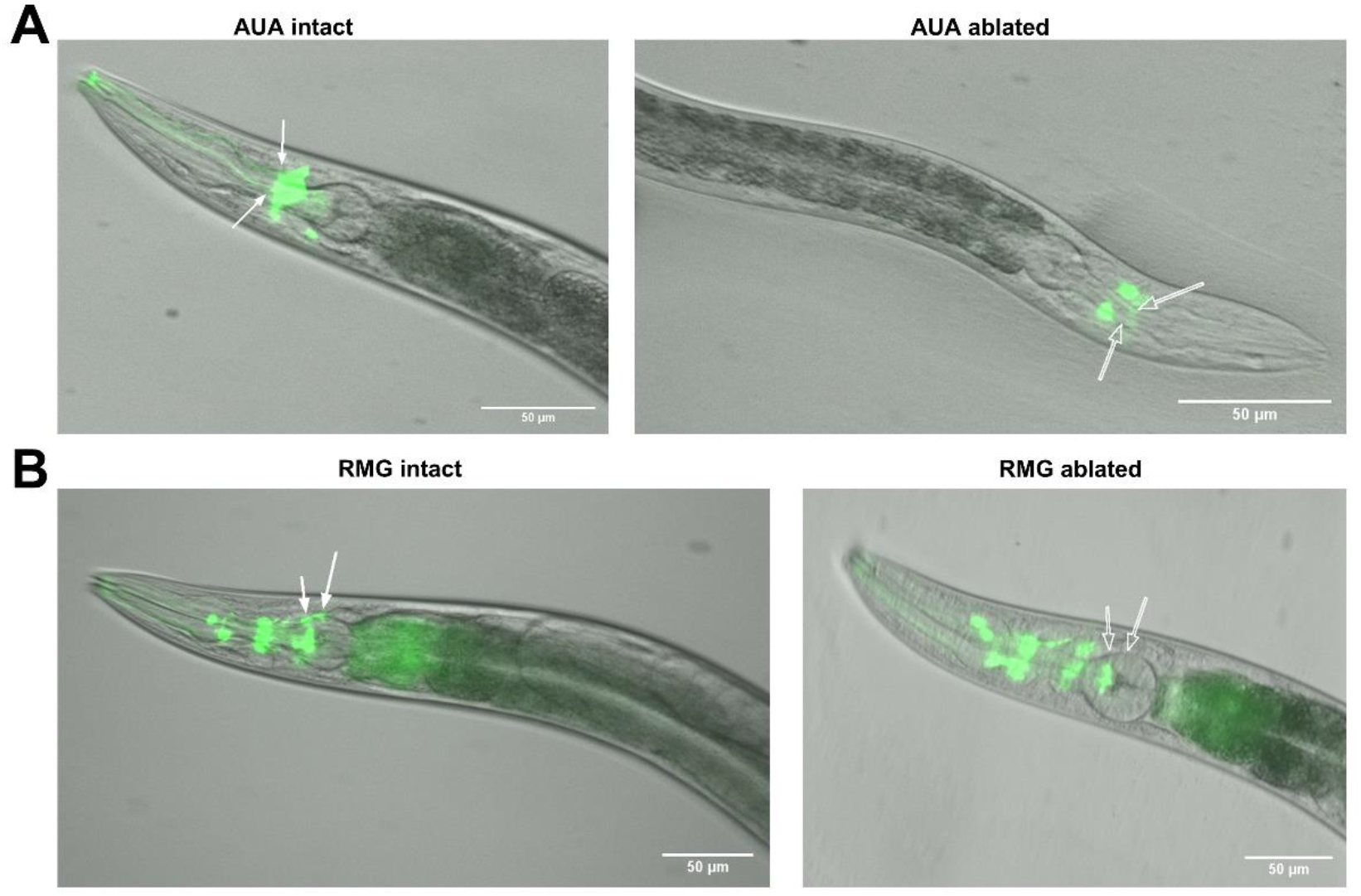
Genetic ablation of AUA and RMG neurons. (A) Representative fluorescent micrographs of NY2078 ynIs78 [*flp-8p::GFP*] (left) and AY178 ynIs78 [*flp-8p::GFP*]; *flp-8p::ced-3 (p15)::nz + flp-32::cz::ced-3 (p17) + unc-122p::rfp* (right) animals. (B) Representative fluorescent micrographs of NY2087 ynIs87 [*flp-21p::GFP* (left) and AY179 ynIs87 [*flp-21p::GFP*]; *flp-21p::ced-3 (p15)::nz + ncs-1p::cz::ced-3 (p17) + unc-122p::rfp* (right) animals. Greyscale and green fluorescent channels have been merged for all micrographs. White, filled arrows point to intact AUA or RMG neurons, while white, unfilled arrows point to the lack of AUA or RMG neurons. Scale bars are 50 μm.

**Fig. S5.**
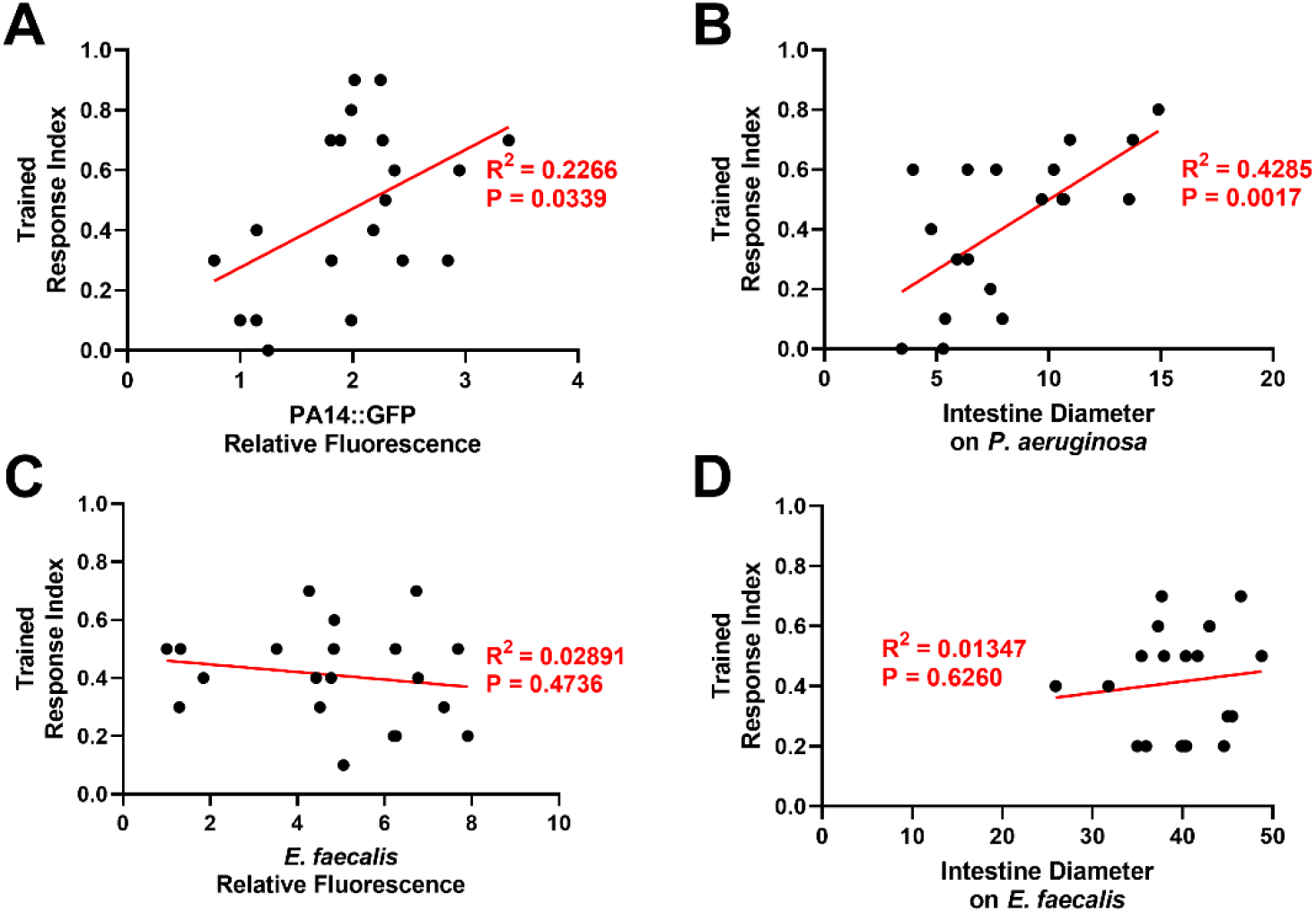
Correlation of intestinal distention and learned reflexive aversion for *P. aeruginosa* but not *E. faecalis* exposure. (A) PA14::GFP relative fluorescence in the intestine (x-axis) and the trained response index to *P. aeruginosa* (y-axis) were measured in individual animals (dots), and linear regression was performed (red line). (B) Same as **A** but with intestinal diameter on *P. aeruginosa* (x-axis). (C) Same as **A** but with *E. faecalis*. (D) Same as **B** but with *E. faecalis*. R^2^ and P-values are shown next to linear regression lines in red.

